# Strand inequality in terms of expression-dependent synonymous single-nucleotide polymorphisms in the *Escherichia coli* chromosome

**DOI:** 10.64898/2026.05.22.727198

**Authors:** Nishita Deka, Pratyush Kumar Beura, Piyali Sen, Ruksana Aziz, Adiraj Kashyap Bora, Deepshikha Keot, Pinju Saikia, Monika Jain, Nima Dondu Namsa, Ramesh Chandra Deka, Edward J Feil, Siddhartha Sankar Satapathy, Suvendra Kumar Ray

## Abstract

Mutations in genomes are mainly attributed to replication, though transcription, which is a more frequent event than replication, is also known to be mutagenic in an organism. However, a distinct demonstration of specific mutations being replication-dependent and/or expression-dependent in genomes is yet to be established. Here, we studied synonymous single-nucleotide polymorphisms (SNPs) in 2091 individual coding sequences (CDS) in the leading strand (LeS) and the lagging strand (LaS) of the *Escherichia coli* chromosome by comparing across 157 strains. The frequencies of complementary transitions (*ti*) and complementary transversions (*tv*) were compared in each CDS to assess parity violation in the context of strand location and gene expression using codon adaptation index (CAI) as a proxy. The C→T and G→A *ti* exhibited the maximum frequency as well as the most prominent strand inequality, as these *ti* were influenced both by the strand asymmetry and gene expression, especially in the LeS. Surprisingly, the inequality between T→C and A→G was found to be gene expression dependent. In addition to CAI-based analysis, we performed the above analysis in CDS based on protein expression data and obtained similar results. A→T and G→T *tv* were more frequent than their complementary T→A and C→A, respectively, in a strand-independent manner. The strand-independent but expression-dependent synonymous SNP inequality in CDS supports the role of transcription-induced mutagenesis contributing to strand inequality in the *E. coli* chromosome.

**Significance Statement:** Mutational patterns in bacterial genomes are shaped by both replication and transcription, but their relative roles remain unclear. By analyzing synonymous SNPs across 157 strains of *E. coli*, a strong asymmetry between the strands regarding C→T and G→A *ti* is observed, as it is influenced by replication asymmetry and expression. We further demonstrate T→C *ti* being enriched in the highly expressed genes and A→G *ti* being enriched in the lowly expressed genes in a strand-independent manner. Notably, replication asymmetry as well as expression influences *ti* SNPs, whereas the relationship between complementary *tv* SNPs are found to be not significantly influenced by expression and strand asymmetry. These findings highlight how some SNPs are influenced by expression but not by replication.

## Introduction

During chromosome replication, the leading strand (LeS) is replicated continuously, while the lagging strand (LaS) is replicated in a discontinuous manner (Okazaki et al. 1968). This asymmetry in replication is already known to cause strand asymmetry in base substitution mutations in genomes (Wu and Maeda 1987), as experimentally demonstrated by Fijalkowska et al (Fijalkowska et al. 1998). The phenomenon of strand inequality is observed more prominently in prokaryotic chromosomes because of the presence of a single origin of replication, unlike in eukaryotic chromosomes (Frank and Lobry 1999). During replication, the LeS remains single-stranded for longer than the LaS, making it more prone to specific chemical changes in DNA, such as cytosine deamination, which gives rise to the most common C→T mutation in the LeS (Lobry and Sueoka 2002). This feature has been exploited to study GC and AT skew in chromosomes, which is used to determine LeS and LaS, along with the origin and terminus of chromosome replication in bacterial chromosomes (Lobry 1996). This is popularly known as parity rule 2 (PR2) or Chargaff’s second parity violation in chromosomes (Sueoka 1995); (Powdel et al. 2009). Transcription also contributes to strand inequality in genomes as non-template DNA temporarily becomes single-stranded, which is more prone to damage such as cytosine deamination and guanine oxidation (Beletskii and Bhagwat 1996). The template strand is better protected due to transcription-coupled repair (TCR) (Francino and Ochman 2001).There is a protein called mfd (*mutation frequency decline*) that acts as a major contributing factor of TC-NCR (Selby et al. 2023). However, Park et al. showed that transcription is overall mutagenic, with higher mutation rates in highly expressed genes in yeast and the human germline. This is due to transcription-associated mutations (TAM), which affect both strands and can outweigh the protective effect of TCR in eukaryotes (Park et al. 2012). Martincorena et al. reported a negative association between gene expression and mutation rate in *Escherichia coli*, with highly expressed genes exhibiting lower mutation rates, potentially reflecting preferential protection or repair of highly expressed genes (Martincorena et al. 2012). Sharp and Li demonstrated a relationship between codon usage bias, gene expression, and substitution (Sharp and Li 1987). The selection-mutation-drift theory suggests that the strength of selection on codon usage bias is different between highly expressed and lowly expressed genes. Selection favors optimal codons for efficient translation, while mutation and drift allow non-optimal codons to persist in the population (Bulmer 1991). Because selection is stronger on highly expressed genes, leading to stronger codon usage bias compared to lowly expressed genes.

The common understanding about C→T enrichment in LeS due to preferred cytosine deamination in single-stranded DNA than double-stranded DNA has been the major explanation in all this research (Lobry 1996),(Frank and Lobry 1999),(Lobry and Sueoka 2002).The same explanation has been given for the coding strands to explain transcription-associated mutation bias (Beletskii and Bhagwat 1996) Research on the chromosomes of different bacteria has sought to elucidate the mechanism underlying PR2 violation or GC skew (Rocha et al. 2006), revealing that different mutations may be responsible for the same skew in chromosomes.

Gene distribution asymmetry between the strands is observed due to the preferential localization of essential and highly expressed genes in the LeS (Rocha and Danchin 2003).In the LeS, both machineries move in the same direction, resulting in co-directional collisions, whereas in the LaS, they move oppositely, leading to head-on collisions that are more detrimental to coding sequences (CDS) and increase mutation rates (Sankar et al. 2016). Srivatsan et al. (2010) demonstrated that head-on transcription–replication conflicts can slow replication speed, disrupt the replication fork, and increase the mutation rate of a gene within an inverted genomic region, suggesting that avoidance of such conflicts may contribute to the evolution of transcription–replication co-orientation (Srivatsan et al. 2010). Consequently, more genes are found in the LeS than in the LaS (Mao et al. 2012), likely to minimize the head-on collision between replication and transcription machineries. Recent studies have investigated how transcription influences mutagenesis in organisms (Adebali et al. 2025),(Carvajal-Garcia et al. 2024). It has been demonstrated that transcription-coupled repair causes mutations through nucleotide excision repair (NER) (Adebali et al. 2017). Oxidative stress inside the cell has been demonstrated to be a major driving force for mutations during transcription, leading to the development of antibiotic resistance in bacteria (Carvajal-Garcia et al. 2023). These studies have opened up an interesting question regarding the possibility of a dominant contribution of transcription to the strand inequality in bacterial chromosomes. Strand inequality might result from the gene distribution asymmetry as well as the difference in the collision between replication and transcription between the strands. The magnitude of contribution of replication asymmetry *per se* towards strand inequality may be lower than that of transcription. It is pertinent to note that a CDS undergoes replication once before cell division, whereas the same CDS might undergo thousands of rounds of transcription with one round of cell division. Therefore, the non-template strand remaining as single-stranded due to transcription is several times higher than the strand remaining single-stranded during replication. Therefore, due to biased gene distribution inequality towards LeS, the amount of time this strand remained single-stranded as a non-template DNA is proportionately more than that of LaS. Therefore, considering transcription-induced mutagenesis, the strand inequality in the genome might be primarily attributed to transcription than replication, and to preferred gene location to LeS.

Despite a plethora of research on strand inequality in terms of mutation attributing to replication asymmetry and transcription, a distinct understanding of replication and/or expression influencing certain *tis* and *tv*s out of the twelve possible SNPs has not been adequately reported in any of this research. A proportional demonstration of gene expression and frequency of certain *ti* or *tv* will be important to address the above queries. The genome sequence of many strains belonging to a species is available now. These sequences can be analyzed to identify synonymous variation in genes, which is under weak purifying selection. Therefore, these variations are likely to represent base substitution mutations arising due to replication as well as gene expression. In this study, we have explored the *E. coli* chromosome to study SNPs in the LeS and LaS. Codon adaptation index (CAI) has been used to identify highly expressed and lowly expressed genes to study the impact of gene expression on SNPs.CAI is a highly accepted proxy for gene expression in case of *E. coli* and can be calculated for nearly all protein-coding genes; it has been shown to correlate strongly with protein abundance in the case of *E. coli* (r = 0.7-0.8). However, experimental conditions, including growth conditions and environmental factors, often influence protein-level expression (Schmidt et al., 2016; Mori et al., 2021). Considering the well-established role of codon usage in determining translational efficiency, particularly in rapidly growing bacteria such as *E. coli*, CAI was therefore primarily used as a consistent proxy for relative gene expression potential in the present study. Nevertheless, to assess the robustness of the observed trends, the analysis was also performed using experimentally determined protein abundance values. Importantly, both approaches yielded comparable patterns, providing additional support for the reliability of the CAI-based observations.

Apart from the role of replication in C→T and G→A, an influence of gene expression on this variation is observed in this study. Interestingly, variation such as T→C and A→G has been observed to be increased by highly and lowly expressed genes, respectively. Variations such as A→T and G→T have been observed to be influenced by gene expression but not by the strand asymmetry. Overall, the distinct roles of replication and gene expression on different SNPs has been observed in this study, which demonstrates the effect of gene expression being confounded with replication while studying strand asymmetry in bacterial chromosomes.

## Result

### Reverse trend of complementary transition SNPs between the strands

Synonymous SNPs were analyzed in the CDS of the LeS and the LaS separately in *E. coli* chromosome to investigate the strand-specific bias. Of the 2091 CDS filtered for the analysis, 1166 were from the LeS and 925 were from the LaS. The maximum size CDS was 4554 bp, whereas the minimum size CDS was 195 bp. There was no strand bias observed regarding CDS size. The abundance values of SNPs were proportional to the size of the CDS. To understand whether the analysis was not affected by limited abundance values of SNPs, the analysis was performed across CDS of different size categories: (i) all CDS; (ii) ≥ 500 bp; (iii) ≥ 1000 bp; and ≥ 2000 bp. The results discussed below were limited to observations in CDS ≥ 2000 bp (Table 1), though studies were done in all CDS. The frequencies of all twelve types of SNPs across the different CDS groups are presented in Table S1 (supplementary Table 1).

**Table 1:**
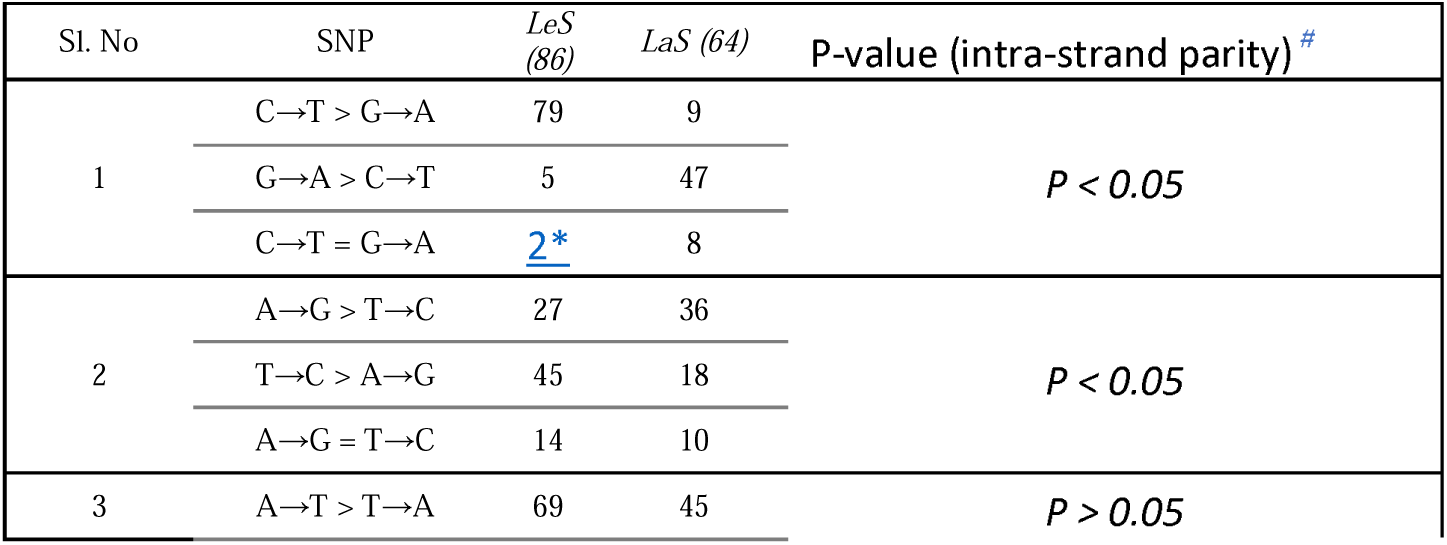
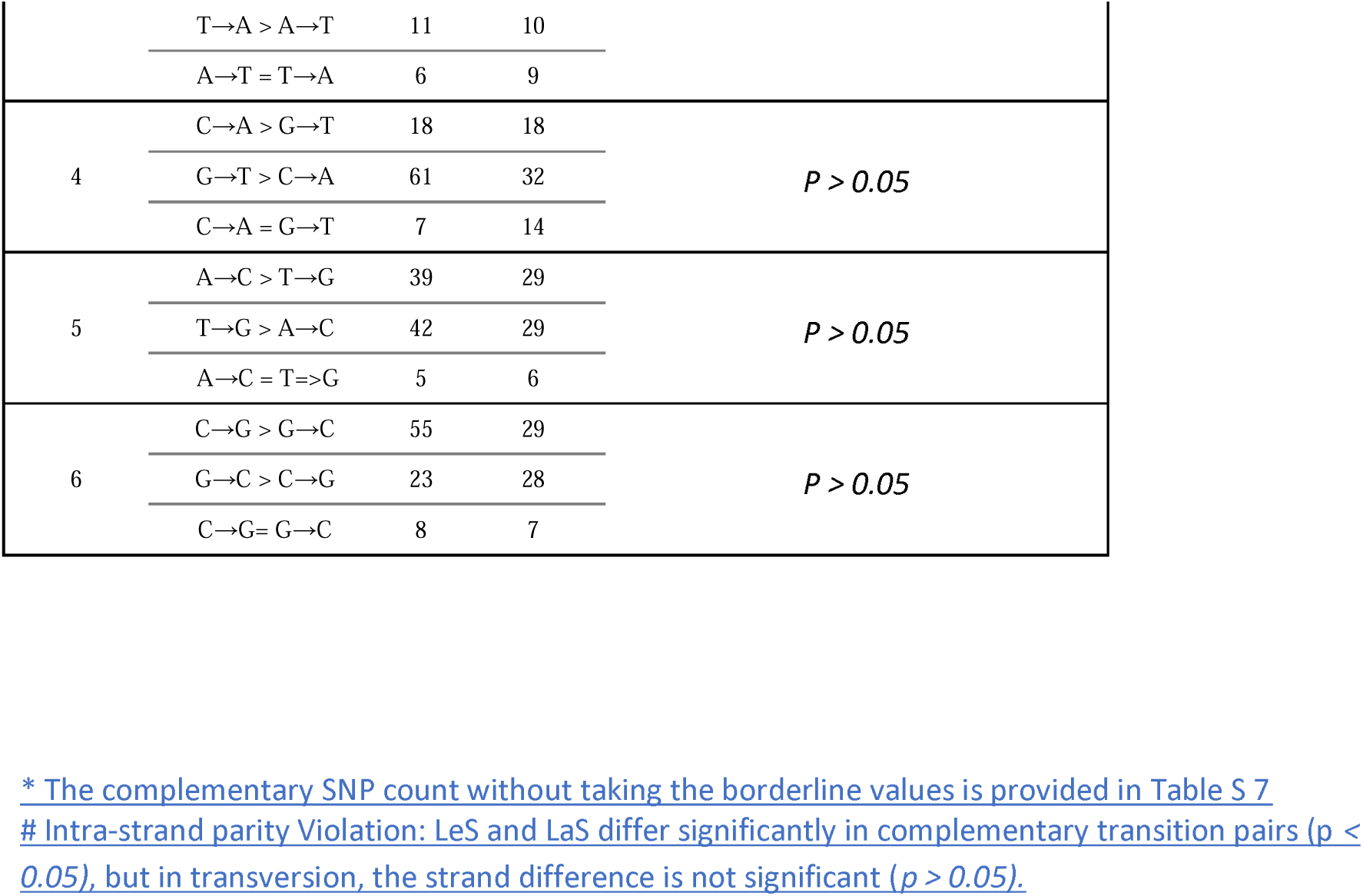
Comparative analysis of complementary SNPs between LeS and LaS. The *ti* complementary pairs are found to be significantly influenced by strand asymmetry (Pearson chi-square test, *p <* 0.05). The *tv* complementary pairs are found to be unaffected by strand asymmetry *p* > 0.05)

The four transitions (*ti*s) exhibited general differences among each other regarding their frequencies: the complementary pair, such as C→T and G→A, were more frequent than the other complementary pair, A→G and T→C, in both strands. A comparative analysis was performed on the frequencies of complementary *ti*s, within a CDS. Among the CDS of ≥ 2000 bp, 86 were in the LeS, and 64 were in the LaS. In the case of LeS, 79 CDS (91.86%) exhibited C→T > G→A, only 5 CDS (5.8%) exhibited the reverse, and 2 CDS exhibited similar frequencies. In the LaS, 47 CDS (73.44%) exhibited G→A > C→T, only 9 CDS (14.06%) exhibited the reverse, and 8 CDS exhibited similar frequencies. There was a distinct reverse pattern was observed between the strands regarding C→T *vs* G→A (Fig.1.1). The same reverse pattern between the strands was also observed in CDS with sizes ≥ 1000 and ≥ 500 bp (Fig. S1). This indicated that the C→T and G→A frequency bias between the strands was strong (*p* < 0.05) (Table S2). The magnitude of PR2 violation was more prominent in larger CDS than in smaller CDS. This might be due to the greater abundance of SNPs in larger CDS than in smaller CDS. However, the possible role of any selection due to the size of CDS cannot be ruled out.

**Fig. 1.1.**
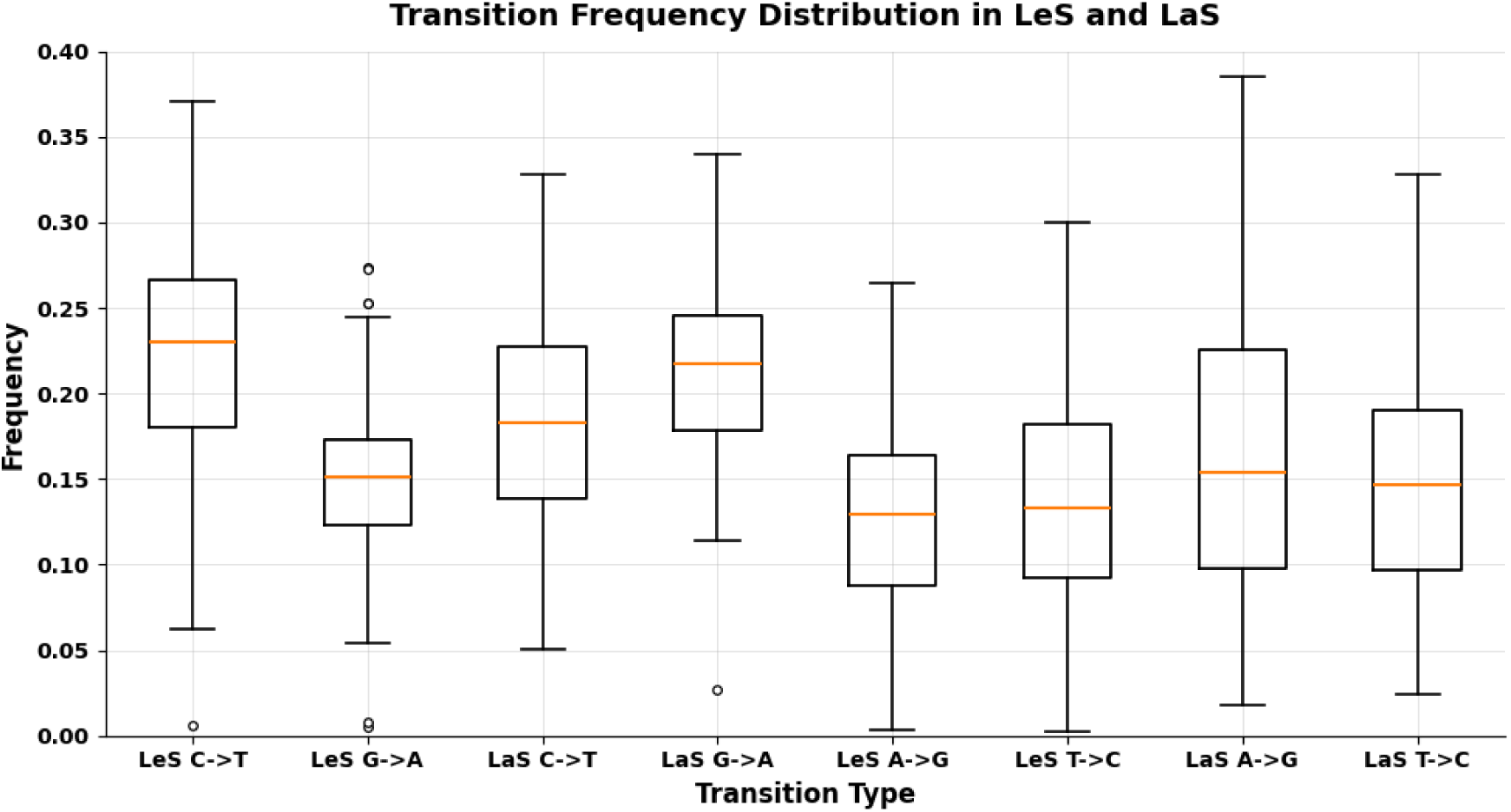
Comparative transition frequency boxplot in both LeS and LaS CDS. Transition frequency of CDSs larger than 2000 bp in both LeS and LaS. In LeS, the frequency of C→T is significantly greater than that of G→A, whereas in LaS the trend is opposite. The frequencies of A→G and T→C have also shown bias towards LaS and LeS, respectively, but it is not found to be statistically significant.

In the case of the other *ti*s such as T→C and A→G, in the LeS, 46 CDS (53.49%) exhibited T→C > A→G, 27 CDS (31.40%) exhibited the reverse, and 13 CDS exhibited similar frequencies. In the LaS, 36 CDS (56.25%) exhibited A→G > T→C, 19 CDS (29.69%) exhibited the reverse, and 10 CDS exhibited similar frequencies. This indicated a lower but consistent T→C bias in the LeS and an A→G bias in the LaS (Fig. 1.1), which were also observed across other CDS of different sizes (≥ 500 bp and ≥ 1000 bp). The A→G and T→C (frequency) strand-specific violations were not significant (*p* > 0.05), unlike C→T vs G→A in all CDS groups (Table S2). It is pertinent to note that in both complementary pairs of *tis*, the bias was more prominent in the LeS than in the LaS.

We then compared the frequencies of each *ti* between the two strands. The average C→T frequency was 0.22 in the LeS and 0.18 in the LaS (*p* < 0.05), whereas the average G→A frequency was 0.15 in the LeS and 0.21 in the LaS (*p* < 0.05). This was in concordance with the strand bias observed above. Similarly, the average A→G frequency was 0.12 in the LeS and 0.16 in the LaS (*p* < 0.05), whereas the average T→C frequency was 0.13 in the LeS and 0.14 in the LaS (*p* > 0.05). The frequency values between the strands indicate enrichment of T nucleotides and depletion of C nucleotides in the LeS. In the LaS, G→A frequency was higher than that of A→G, which indicated G nucleotide depletion in this strand. Except for C→T, frequency values of the other three *ti* changes were higher in the LaS than in the LeS. The dynamics between Pu→Pu were higher in the LaS (0.38) than in the LeS (0.28), and the dynamics of Py→Py were higher in the LeS (0.36) than in the LaS (0.33). The *ti* frequency in LeS was 0.64, whereas that in LaS was 0.71, indicating that the overall *ti* frequency was higher in LaS than in LeS. The *ti* frequency in both strands was biased toward A/T.

### Complementary transversion SNPs exhibit a similar pattern between the strands

The frequencies of eight single-nucleotide *tvs*, such as A→C, A→T, C→A, C→G, G→C, G→T, T→A, and T→G, were found in the CDS of both strands. Frequencies were different for different *tv*s. A→T frequency was highest among all tvs, whereas G→C frequency was lowest. An intra-strand comparison of complementary *tv*s for CDS sizes ≥ 2000 bp was performed (Fig. 1.2). In the LeS, out of 86 CDS, 69 (80.23%) exhibited A→T >T→A, 11 CDS (12.79%) exhibited the reverse, and 6 CDS exhibited similar frequencies. In the LaS, out of 64 CDS, 45 (70.31%) exhibited A→T >T→A, 10 CDS (15.63%) exhibited the reverse, and 9 CDS exhibited similar frequencies. This indicated that the two strands exhibited a similar pattern regarding A→T and T→A. In another complementary pair, 61 CDS (70.93%) in the LeS and 32 CDS (50.00%) in the LaS exhibited G→T > C→A. Thus, in the above two complementary pairs, the trend was strand-independent. Overall, A→T and G→T substitutions occurred more frequently than their complementary counterparts T→A and C→A in both strands. PR2 violation was observed in both strands (*p* < 0.05) (Table S2). This pattern was consistent across CDS of sizes ≥500 bp and ≥1000 bp. In the LeS, C→G > G→C was observed in 55 CDS (63.95%), and G→C > C→G was observed in 23 CDS (26.74%), and 8 exhibited G→C = C→G, whereas in the LaS, C→G > G→C was observed in 29 CDS (45.31%), and G→C > C→G was observed in 28 CDS (43.75 %), and 7 exhibited G→C = C→G. The bias was observed only in the LeS with PR2 violation (*p* < 0.05) (Table S2). This pattern was consistent across CDS of ≥1000 bp and ≥500 bp. In the LeS, T→G > A→C was observed in 42 CDS (48.84%), whereas A→C > T→G was observed in 39 CDS (45.35%), and 5 CDS exhibited the same frequencies. In the LaS, A→C > T→G was observed in 29 CDS (45.31%), whereas T→G > A→C was observed in 29 CDS (45.31%), and 6 CDS exhibited the same frequency. The PR2 rule was maintained in both strands (*p* > 0.05) (Table S2). It was observed that complementary *tv*s exhibited the same trend in LeS and LaS, unlike complementary *ti*s, where strand bias is stronger (Fig. 1.1).

**Fig. 1.2:**
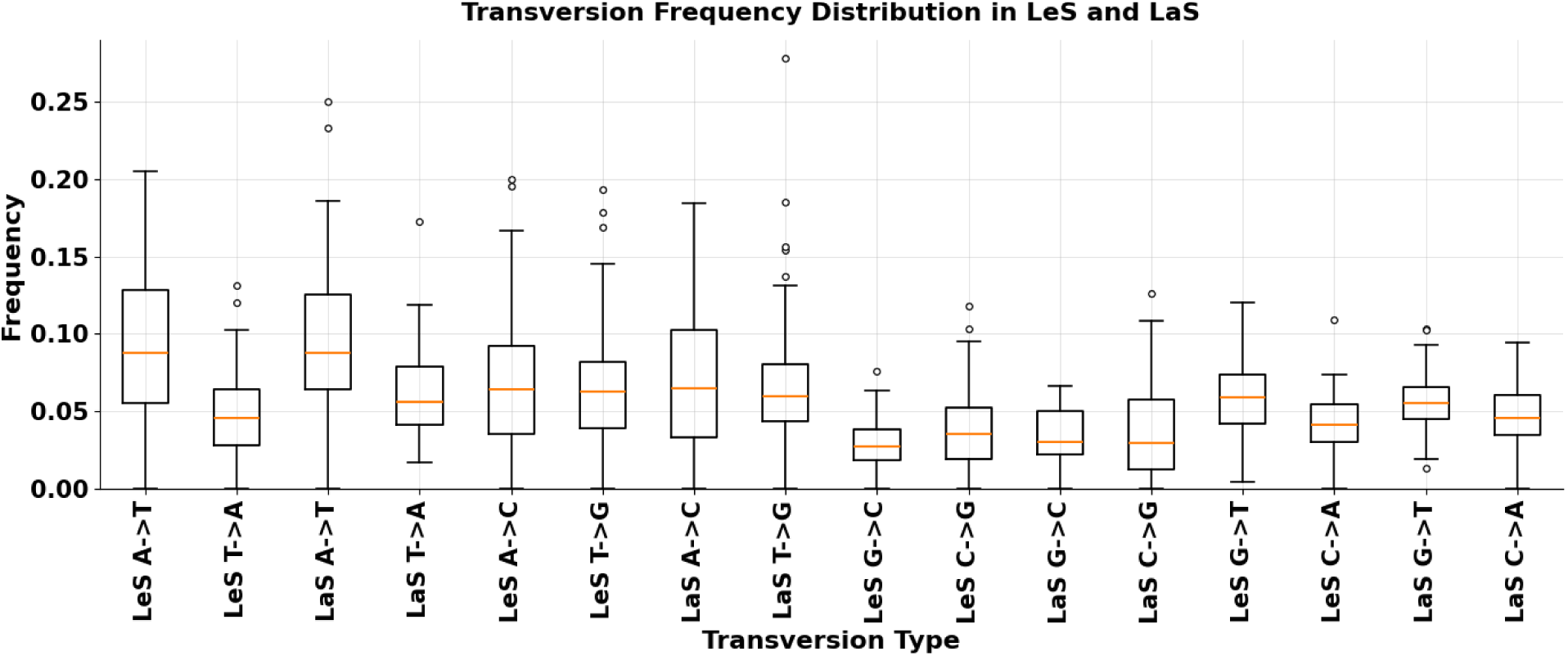
Transversion frequency of CDSs larger than 2000 bp. Transversion frequency of CDSs larger than 2000 bp in both LeS and LaS. A→T frequency was the maximum among all *tv* types, whereas G→C frequency was the minimum. Unlike *ti* complementary pair, the transversion complementary pairs show no strand dependency. This indicated that the *ti* frequency was more strand-dependent, which was statistically proven using Pearson Chi-Square statistics (Table 1)

We compared the frequencies of each *tv* between the strands. In CDS ≥ 2000 bp, the average frequency of A→T was 0.092 in the LeS and 0.096 in the LaS, whereas the average frequency of T→A was 0.048 in the LeS and 0.061 in the LaS. The G→T frequency was 0.058 in the LeS and 0.056 in the LaS, whereas the C→A frequency was 0.042 in the LeS and 0.046 in the LaS. A→C frequency was 0.067 in the LeS and 0.070 in the LaS, whereas the T→G frequency was 0.064 in the LeS and 0.067 in the LaS. The G→C frequency was 0.028 in the LeS and 0.030 in the LaS, whereas the C→G frequency was 0.037 in the LeS and 0.036 in the LaS. These observations were in concordance with the intra-strand comparison of *tv* frequencies. Except for T→A (*P < 0.05*), which was more frequent in the LaS than in the LeS, no other *tv* was significantly different between the strands. Overall, the frequency of *tv* is higher in the LaS (0.46) than that in the LeS (0.43), like *ti*. In the LeS, Pu→Py dynamics (0.24) were higher than Py→Pu dynamics (0.19). Likewise, in the LaS, Pu→Py dynamics (0.26) were higher than Py→Pu dynamics (0.21). These observations suggest a consistent bias toward Pu→Py substitutions over Py→Pu substitutions in both strands. Inter-strand mutation comparison data for CDS ≥1000 and ≥500 are given in Table S1. Our results indicated that there was no replication-induced bias regarding transversion between the strands.

### Influence of gene expression on transition SNPs

It is already known that the LeS has proportionately more highly expressed genes than lowly expressed genes compared to LaS (Rocha and Danchin 2003). In this study, we observed that the frequency distribution between complementary pairs was more biased in LeS than in LaS. So, we studied the influence of gene expression on SNPs. Codon Adaptation Index (CAI) can be used as an expression index in *E. coli*, as CAI is proportional to gene expression (Sharp and Li 1987). For CDS ≥ 1000 bp, the average CAI value of the top 50 CDS with the maximum CAI values in LeS and LaS was 0.612 and 0.547, respectively. The average CAI value of the 50 CDS with the minimum CAI values in LeS and LaS was 0.246 and 0.247, respectively. The boxplot diagram of highly and lowly expressed genes is shown in Fig. 2. It was consistent with established studies that highly expressed genes were more dominant in LeS than in LaS. The 50 CDS with maximum CAI values in both strands were referred to as highly expressed genes, whereas the 50 CDS with minimum CAI values in both strands were referred to as lowly expressed genes.

**Fig. 2:**
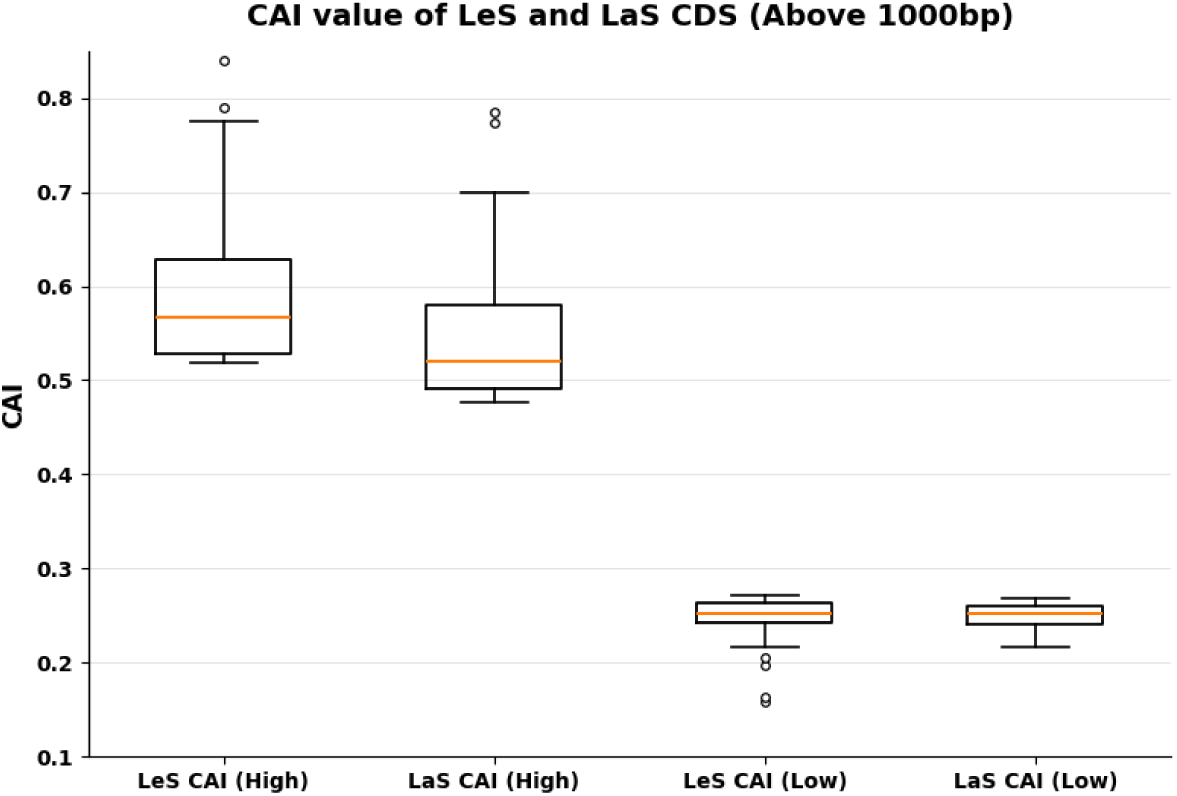
Comparison of the expression levels between LeS and LaS CDS. Distribution of highly expressed genes and lowly expressed genes across LeS and LaS strands. The distribution of HEGs is uneven *(P <* 0.05), with a predominant presence in LeS compared to LaS. In contrast, lowly expressed genes show no significant difference in distribution between the two strands *(P >* 0.05).

We studied C→T and G→A transitions in both highly and lowly expressed genes on both strands. In the LeS, out of 50 highly expressed CDS, 48 exhibited C→T > G→A, and only 1 exhibited the reverse, and 1 exhibited equal frequency, whereas out of the 50 lowly expressed CDS, 37 exhibited C→T > G→A, 9 exhibited the reverse, and 4 exhibited equal frequency. In the LaS, out of the top 50 highly expressed CDS, 27 exhibited G→A > C→T, 15 exhibited the reverse, and 8 exhibited equal frequency; whereas, out of the 50 lowly expressed CDS, 34 exhibited G→A > C→T, 9 exhibited the reverse, and 7 exhibited equal frequency.

In LeS, the C→T frequency was 0.216 and 0.203 in highly expressed genes and lowly expressed genes, respectively. In LaS, the C→T frequency was 0.154 in highly expressed genes and 0.146 in lowly expressed genes, respectively. The gradual decrease in frequency from LeS to LaS, as well as from highly expressed genes to lowly expressed genes, was distinctly observed. In LeS, the G→A frequency was 0.136 in highly expressed genes and 0.155 in lowly expressed genes, respectively. In LaS, the G→A frequency was 0.167 and 0.183 in highly expressed genes and lowly expressed genes, respectively. The gradual increase in frequency from the LeS to the LaS and from the highly expressed genes to the lowly expressed genes was distinctly observed. (Table 2.2). The impact of strand location is prominent as is the influence of expression (Table 2.1).

**Table 2.1.** Effect of gene expression on complementary SNPs in both LeS and LaS. The complementary SNPs were quantified to analyze the effect of gene expression on both strands. The A→G and T→C frequencies were found to be highly influenced by gene expression in both strands. The C→T and G→A complementary *ti,* and G→T and C→A complementary *tv* were influenced by gene expression in case of LeS but not in LaS. A→T and T→A complementary tv is influenced by gene expression in the case of LaS but not in LeS.

| **Sl. no** | **Complementary Pair** | **HEG_LeS** | **LEG_LeS** | ***p* value #** | **HEG_LaS** | **LEG_LaS** | ***p*-value #** |
|---|---|---|---|---|---|---|---|
| **1** | C→T > G→A | **48** | **37** | ***P* < 0.05** | **15** | **9** | ***P* > 0.05** |
|  | G→A > C→T | **1** | **9** |  | **27** | **34** |  |
|  | C→T = G→A | **1** | **4** |  | **8** | **7** |  |
| **2** | A→G > T→C | **8** | **32** | ***P* < 0.05** | **15** | **34** | ***p* < 0.05** |
|  | T→C > A→G | **38** | **15** |  | **30** | **13** |  |
|  | T→C = A→G | **4** | **3** |  | **5** | **3** |  |
| **3** | A→C > T→G | **24** | **20** | ***P* > 0.05** | **28** | **24** | ***p* > 0.05** |
|  | T→G > A→C | **24** | **25** |  | **19** | **22** |  |
|  | T→G = A→C | **2** | **5** |  | **3** | **4** |  |
| **4** | A→T > T→A | **40** | **32** | ***P* > 0.05** | **37** | **28** | ***p* < 0.05** |
|  | T→A > A→T | **9** | **17** |  | **9** | **20** |  |
|  | T→A = A→T | **1** | **1** |  | **4** | **2** |  |
| **5** | G→T > C→A | **37** | **25** | ***P* < 0.05** | **26** | **31** | ***p* > 0.05** |
|  | C→A > G→T | **11** | **20** |  | **20** | **17** |  |
|  | C→A =G→T | **2** | **5** |  | **4** | **2** |  |
| **6** | C→G > G→C | **26** | **28** | ***p* > 0.05** | **27** | **21** | ***p* > 0.05** |
|  | G→C > C→G | **17** | **19** |  | **15** | **24** |  |
|  | G→C = C→G | 7 | 3 |  | **8** | **5** |  |

\# *p*-values of chi-square statistics were shown. The complementary SNPs are compared with respect to gene expression in both LeS and LaS

**Table 2.2:** Frequency of complementary SNPs in HEG and LEG genes of LeS and LaS. Frequencies of all 12 types of substitution for the CDS of LeS and LaS above 1000 bp are shown. The C→T and G→A frequencies were strand-dependent and high in LeS and LaS CDS, respectively. The frequencies of A→G and T→C are found to be expression-dependent, where A→G frequency was higher than T→C in LEG, and vice versa was true for HEG genes.

| **Strand** | **Status** | **C→T** | **G→A** | **A→G** | **T→C** | **A→T** | **T→A** | **C→A** | **G→T** | **C→G** | **G→C** | **A→C** | **T→G** |
|---|---|---|---|---|---|---|---|---|---|---|---|---|---|
| **LeS** | All CDS (≥ 1000 bp) | 0.223 | 0.150 | 0.125 | 0.131 | 0.088 | 0.051 | 0.041 | 0.059 | 0.035 | 0.029 | 0.060 | 0.064 |
|  | All CDS (≥ 2000 bp) | 0.223 | 0.150 | 0.130 | 0.140 | 0.092 | 0.048 | 0.043 | 0.059 | 0.038 | 0.028 | 0.067 | 0.065 |
|  | HEG (≥ 1000 bp) | 0.216 | 0.136 | 0.100 | 0.139 | 0.102 | 0.048 | 0.036 | 0.062 | 0.032 | 0.025 | 0.051 | 0.054 |
|  | LEG (≥ 1000 bp) | 0.203 | 0.155 | 0.120 | 0.092 | 0.074 | 0.049 | 0.042 | 0.051 | 0.032 | 0.026 | 0.053 | 0.055 |
| **LaS** | All CDS ≥ 1000 bp | 0.176 | 0.201 | 0.140 | 0.129 | 0.091 | 0.052 | 0.045 | 0.045 | 0.031 | 0.032 | 0.064 | 0.058 |
|  | All CDS ≥ 2000 bp | 0.186 | 0.214 | 0.163 | 0.148 | 0.097 | 0.061 | 0.047 | 0.056 | 0.037 | 0.034 | 0.070 | 0.067 |
|  | HEG (≥ 1000 bp) | 0.154 | 0.167 | 0.101 | 0.115 | 0.088 | 0.043 | 0.035 | 0.047 | 0.026 | 0.021 | 0.046 | 0.035 |
|  | LEG (≥ 1000 bp) | 0.146 | 0.183 | 0.117 | 0.085 | 0.064 | 0.051 | 0.043 | 0.053 | 0.025 | 0.027 | 0.047 | 0.046 |

We studied A→G and T→C transitions in both highly and lowly expressed genes on both strands. In the LeS, out of the 50 highly expressed CDS, 38 exhibited T→C >A→G, 8 exhibited the reverse, and 4 exhibited equal frequency; whereas, out of the 50 lowly expressed CDS, 15 exhibited T→C >A→G, 32 exhibited the reverse, and 3 exhibited equal frequency. In the LaS, out of the 50 highly expressed CDS, 30 exhibited T→C >A→G, 15 exhibited the reverse, and 5 exhibited equal frequency, whereas out of the 50 lowly expressed CDS, 13 exhibited T→C >A→G, 34 exhibited the opposite pattern, and 3 exhibited equal frequency. Pearson chi-square test <u>was also performed to obtain the statistical significance of the data</u>, where it was confirmed that the frequency of A→G and T→C was significantly influenced by gene expression in both strands (*p* < 0.05) (Table 2.1). This <u>study was also performed using protein expression data and obtained the similar result (Table S8)</u>

In LeS, the A→G frequency was 0.100 in highly expressed genes and 0.120 in lowly expressed genes. In LaS, the corresponding frequencies were 0.101 and 0.117, respectively. Overall, the A→G frequency was higher in lowly expressed genes than in highly expressed genes in both strands. In LeS, the T→C frequency was 0.139 in highly expressed genes and 0.092 in lowly expressed genes. In LaS, the corresponding frequencies were 0.115 and 0.085, respectively. Thus, the T→C frequency was consistently higher in highly expressed genes than in lowly expressed genes in both strands (Fig. 3.1). The T→C frequency was higher in LeS than in LaS and higher in highly expressed genes than in lowly expressed genes. The frequency of A→G was lower in LeS than in LaS and lower in highly expressed genes than in lowly expressed genes (Fig 3.2). The impact of expression was prominent, while the influence of strand location was mild on these variations. Another observation regarding *Sti* was that, in highly expressed genes, Py→Py (0.624) variations were more than Pu→Pu (0.504) variations, while in the lowly expressed genes, Pu→Pu (0.575) variations were more than Py→Py (0.526).

**Fig 3.1:**
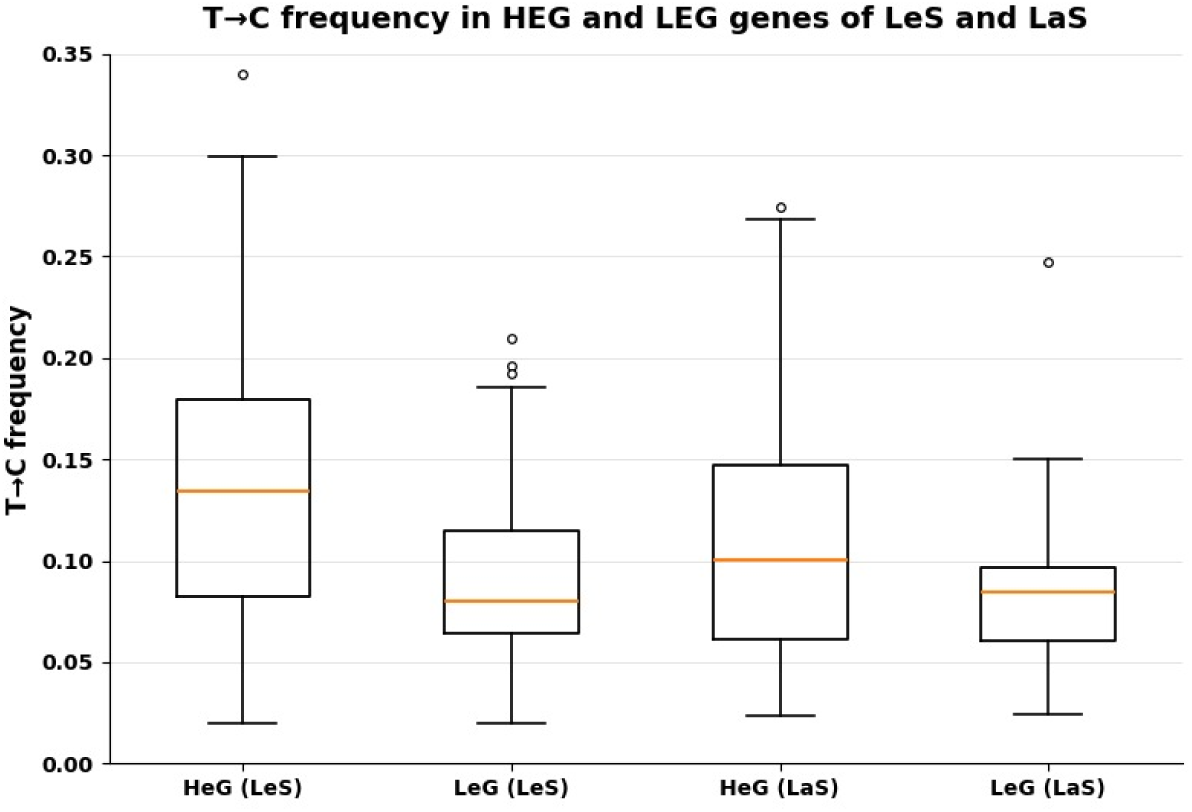
Comparison of T→C frequency between highly expressed genes and lowly expressed genes and LaS and LeS. The frequency of T→C was compared between the highly expressed and lowly expressed genes of both strands, and it was found that the T→C frequency was higher in highly expressed genes than in lowly expressed genes in both LeS and LaS

**Fig. 3.2:**
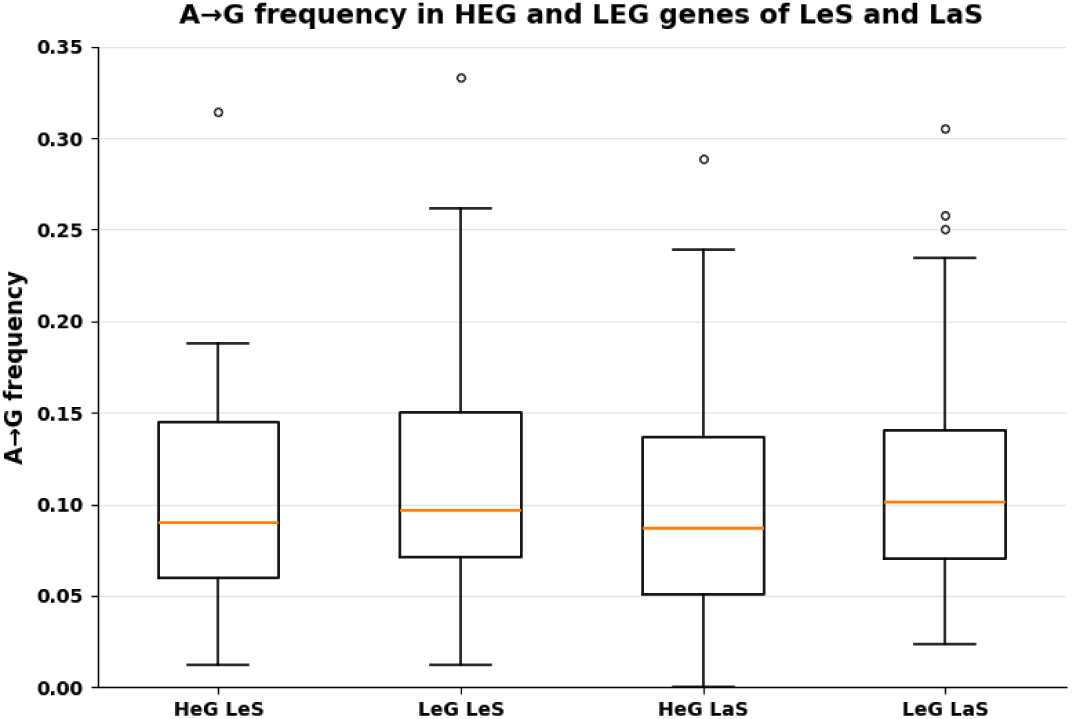
Comparison of A→G frequency between highly expressed genes and lowly expressed genes and LaS and LeS.

Our previous analysis regarding replication-associated *ti*s indicated that the LeS exhibited bias toward T→C, whereas the LaS exhibited bias toward A→G, but subsequent expression analysis suggested that earlier interpretation regarding strand asymmetry was inappropriate. The trend was more pronounced due to the predominance of highly expressed genes on the LeS than on the LaS. This is an example of specific *ti* bias in genes influenced by their expression but not by strand expression.

### Influence of gene expression on *tv*

We further analyzed the influence of gene expression on transversion. The A→T frequency was higher than the T→A frequency in both the LeS and LaS, irrespective of gene expression level. Although the A→T frequency was influenced by gene expression, the T→A frequency showed comparatively little variation with gene expression. The influence of gene expression on this complementary *tv* was found to be statistically significant in LaS CDS (Pearson chi-square statistics, *p* < 0.05) (Table 2.1).

The G→T frequency was higher than C→A in highly expressed genes than in lowly expressed genes in the LeS (Pearson chi-square statistics, *p* < 0.05) (Table 2.1) but not in LaS. Other than these two complementary pairs, no other transversion pairs were found to be influenced by gene expression in any of the strands. The quantification of complementary transversions is presented in Table 2.1, while their corresponding frequencies are presented in Table 2.2. Since *tv* frequencies were not significantly affected by gene expression, their results are discussed less extensively than those of transitions.

### High frequency of *ti* over *tv* in the strands

We compared *ti* and *tv* abundances to provide several evidences in support the findings based on consensus sequence–based approach in this study. Synonymous *ti* was found to occur more frequently than synonymous *tv* in both strands. For CDS ≥ 2000 bp, the overall *ti/tv* ratio was 3.42 in the LeS and 3.46 in the LaS. The *ti/tv* ratios were also calculated for different polymorphisms (Table 3). In the LeS, the highest *ti/tv* ratio, 5.95, was observed for C→T/C→G substitutions, and the lowest, 1.41, was observed for A→G/A→T. In the LaS, the highest *ti/tv* ratio was 6.27 observed for G→A/G→C substitutions, and the lowest, 1.68 was observed for A→G/A→T. Codon-wise, SNPs were significantly higher than the double- or triple-nucleotide polymorphisms. Among CDS larger than 2000 bp, approximately 95% were SNPs, whereas the rest were ∼5% double-nucleotide polymorphisms and ∼0.3% triple-nucleotide polymorphisms (Table S3). This pattern was consistent in both LeS and LaS. In the cases of double-nucleotide polymorphisms at a single codon position (single-site double mutations), mostly in the FFD codons, the mutations were predominantly *ti*–*tv* rather than *tv*–*tv*. Our results showed that approximately 80% of such double mutations consisted of one *ti* and one *tv*, indicating a strong preference for *ti–tv* combinations over *tv–tv* combinations. Lastly, synonymous mutations were more abundant than non-synonymous mutations. Though we have elaborately discussed the synonymous part in this study, our analysis revealed a significantly higher synonymous-to-non-synonymous mutation ratio. In this study, the synonymous SNP analysis in 2091 CDS of *E. coli* chromosome was performed using a consensus sequence–based approach, which corroborated several fundamental characteristics of base substitution mutations.

**Table 3:** *Sti/Stv* ratio of individual substitutions is shown. The overall *ti/tv* ratio was 3.42 in the LeS and 3.46 in the LaS. In the LeS, the highest ti/tv ratio, 5.95, was observed for C→T/C→G substitutions, and the lowest, 1.41, was observed for A→G/A→T. In the LaS, the highest ti/tv ratio was 6.27 observed for G→A/G→C substitutions, and the lowest, 1.68, was observed for A→G/A→T.

| *sti/stv* | **LaS** | **LeS** |
|---|---|---|
| **A→G freq/A→C freq** | 2.32 | 1.928 |
| **A→G freq/A→T freq** | 1.684 | 1.41 |
| **C→T freq/C→A freq** | 3.964 | 5.243 |
| **C→T freq/C→G freq** | 5.053 | 5.954 |
| **G→A freq/G→C freq** | 6.271 | 5.279 |
| **G→A freq/G→T freq** | 3.818 | 2.563 |
| **T→C freq/T→A freq** | 2.404 | 2.884 |
| **T→C freq/T→G freq** | 2.19 | 2.151 |
| **Average** | **3.463** | **3.427** |

## Discussion

Strand inequality in chromosomes is well established in the literature, which is attributed to both replication and transcription (Lobry and Sueoka 2002). However, replication asymmetry has been thought of as the main contributor to it, even though it is known that a genomic region undergoes many more rounds of transcription than replication for a given cell cycle. Therefore, the genomic region temporarily remains single-stranded more often due to transcription than replication. Considering this, it can be argued that damage in the genome due to the temporary formation of a single-stranded region occurs more frequently due to transcription than replication. Recently, there have been several reports regarding transcription-induced mutagenesis in organisms (Fousteri and Mullenders 2008; Hanawalt and Spivak 2008; Adebali et al. 2017; Daniel et al. 2018; Carvajal-Garcia et al. 2023; Carvajal-Garcia et al. 2024). In this context, understanding the distinct attributes of replication and transcription towards strand inequality in chromosomes is important.

In this study, synonymous SNPs have been analyzed in 2091 individual CDS in the LeS and LaS of the *E. coli* chromosome to understand the impact of replication (Whether the gene is located on the LeS or LaS) and gene expression regarding the inequality between complementary *ti* and *tv* in the CDS. Considering weak purifying selection on the synonymous SNPs, it is expected that the SNP spectra will be close to the naturally occurring mutations patterns during replication and transcription. C→T and G→A are the most frequent SNPs and exhibit the most obvious inequality between them according to the strands because these substitution mutations are influenced by both replication and gene expression. It is known that genes are not only preferably located in the LeS than the LaS, but the highly expressed genes are also preferably located in the LeS than the LaS (Rocha and Danchin 2003). This is an example where the combined effects of replication-specific bias and expression-specific bias cumulatively shape the strand inequality. The cytosine deamination hypothesis for single-stranded DNA during replication and transcription can explain well the strong bias between the strands regarding C→T and G→A (Georgakopoulos-Soares et al. 2021). Understanding the impact of gene expression and replication on SNPs may help us in the future to understand the evolutionary pattern of the coding sequences based on their strand location, as well as expression. It is pertinent to note that the *ti* complementary pairs are showing strand dependency (*p* < 0.05) but not the *tv* pairs (*p* > 0.05) (Table 1).It will be interesting to understand why certain genes exhibit different patterns, being highly expressed or being located in the LeS. For example, out of the 50 highly expressed genes, only 1 gene exhibits C→T < G→A. It may be that the gene was earlier located in the LeS, but now it is located in the LaS due to inversion. An additional explanation may be that this is likely a lowly expressed gene. However, gene inversions in the populations were not accounted for while interpreting the strand-specific patterns. Assigning every gene to a gene inversion event for LeS and LaS is a huge task, and there are no such methods available for doing it. The distribution of genes is mostly strand-specific, and essential and highly expressed genes are predominantly present in LeS throughout evolution (Rocha 2004). For example, almost 85% of ribosomal genes of any bacterium are predominantly found on the leading strand and near the OriC, specifically in the case of the rapid growing bacteria such as *E. coli*. Therefore, we believe gene inversion events in *E. coli* would not affect our mutation analysis significantly.

The frequency of T→C is found to be higher than that of A→G in the highly expressed genes, while A→G frequency is found to be higher than that of T→C in the lowly expressed genes in both strands in our study. The A→G and T→C complementary *ti* exhibits a mild strand bias, which can be explained with the help of the influence of gene expression and the preferred occurrence of highly expressed genes in the LeS. In the case of T→C and A→G, there is a minimal role of replication, but a strong influence of gene expression was observed. This observation is novel in this study. The higher frequency of A→G over T→C in lowly expressed genes may be attributed to the high frequency of adenine deamination, which causes A→G *ti* (Deka et al. 2025). This *ti* is predominantly repaired through the alternate excision repair (AER) pathway (Talhaoui et al. 2013). It may be proposed that in lowly expressed genes, the efficiency of repair mechanisms may not be as robust as in highly expressed genes. Highly expressed genes are more frequently subjected to transcription-induced repair processes (Hanawalt and Spivak 2008),(Fousteri and Mullenders 2008), which enhance the correction of certain types of mutations. In contrast, lowly expressed genes experience comparatively weaker repair activity (Gaillard and Aguilera 2016), potentially allowing specific mutational biases to persist. This differential repair efficiency may therefore contribute to the observed bias toward A→G substitutions over T→C substitutions in lowly expressed genes. It will be interesting to have an empirical demonstration of this *ti* bias due to expression in future research. It may be that a coding sequence exhibiting T→C > A→G has a greater probability of being a highly expressed gene in the genome. Therefore, a substitution pattern might be helpful to predict the gene’s probable expression level (high/low). It is pertinent to note that CUB and CAI are both related to gene expression in *E. coli*. CAI exhibited positive correlation with protein expression data (r = 0.75) and a negative correlation with *Nc* values (r = -0.85). We have also performed the complementary mutation analysis regarding high and low expression levels based on *Nc* and protein expression data and found a similar pattern of complementary SNPs in highly and lowly expressed genes. The complementary mutation analysis data based on *Nc* and protein expression data are provided in Table S8 with Pearson chi-square analysis. The analysis based on protein expression data confirms the expression-based T C and A G *ti* in *E. coli*.

Regarding *tv*, it is observed that the A→T and G→T frequencies are higher than T→A and C→A, respectively, in highly expressed genes than in lowly expressed genes. The intensity of influence of gene expression on these SNPs depends on the strand location. The G→T and C→A differences in highly and lowly expressed genes are significant in the LeS (*p > 0.05*). Whereas, A→T and T→A frequency differences are significant in LaS *(p >0.05*), A→T *tv* is more common in highly expressed genes than lowly expressed genes, indicating a mild influence of gene expression. Recently, Carvajal-Garcia et al. (2023) (Carvajal-Garcia et al. 2023) demonstrated that endogenous oxidative damage to the non-template strand can induce genome-wide mutations via transcription-coupled nucleotide excision repair (TC-NER). Their findings further indicate that oxidative stress drives mutagenesis primarily during transcription rather than replication. However, the study does not elaborate on the specific spectrum of base substitutions generated through TC-NER. In this context, the observed A→T SNPs may arise as a consequence of oxidative lesions processed during transcription-coupled repair. Specifically, A→T mutations occurring on the non-coding (template) strand could represent a protective mechanism, potentially reducing the likelihood of thymidine dimer formation on the coding strand and thereby preserving coding sequence integrity. It remains to be determined how replication and expression increase the frequency of A→T *tv*s in coding regions. The underlying reason for the high A→T *tv* requires further investigation. Higher A→T mutation might also occur via depurination, followed by insertion of an A opposite the abasic site during replication (Park et al. 2012).

It is widely reported in the literature that C→T and G→A substitutions are the most frequent SNPs (Beura et al. 2023). However, our analysis indicates that this pattern is not universally consistent. In highly expressed genes of the LeS, T→C substitutions are sometimes observed more frequently than G→A. Additionally, in certain cases, the frequency of the A→T *tv* exceeds that of some *ti* mutations, which contrasts with the general expectation that *tv*s occur less frequently than *ti*s.

It is well established that transcription-coupled repair (TCR) protects the transcribed strand of DNA by preferentially removing lesions that stall RNA polymerase during transcription (Hanawalt and Spivak 2008). The Mfd protein acts as a link between transcription and DNA repair. Selby et al. explained how Mfd helps *E. coli* repair DNA damage during transcription. When RNA polymerase (RNAP) encounters DNA damage, it got stuck. Mfd recognizes the stalled RNAP and, using ATP energy, changes its shape and moves RNAP away from the damaged DNA. This allows UvrA and UvrB to recognize the damage site. Then UvrC cuts out the damaged DNA, UvrD removes the damaged fragment, and DNA polymerase I and DNA ligase fill and seal the gap (Selby et al. 2023).. However, recent studies have suggested a more complex role for transcription-coupled nucleotide excision repair (TC-NER), (Lindsey-Boltz and Sancar 2021),(Carvajal-Garcia et al. 2023; Carvajal-Garcia et al. 2024). Emerging evidence indicates that the TC-NER process itself can contribute to mutagenesis, particularly under oxidative stress conditions, where mutations are generated during transcription rather than replication. These findings raise an important biological question: Does TC-NER primarily protect transcribed regions from mutations, or can it also promote mutations and thereby contribute to genome evolution?

There are many methods available for mutation analysis; mostly, they are phylogeny reconstruction methods like maximum parsimony and maximum likelihood methods, which are for inter-species mutation analysis whose evolutionary positions are very strict. In these methods, evolutionary positions were known beforehand, and then mutation analysis was done. For intra-species analysis, these methods might not be suitable because intra-species evolution can be parallel and have no evolutionary gaps between each strain; therefore, by these existing methods, we cannot determine exact evolutionary positions of the different strains of the same species like *E. coli*. All the existing methods provide dichotomous phylogenetic trees, but we believe that intra-species phylogenetic trees should be polychotomous (Fig S2), because many species can evolve from many other species in parallel. In the supplementary file (Fig. S3), we have shown 5 genes of *E. coli* for which phylogenetic trees were made using ML methods, and we have found that all the phylogenetic tree patterns are very distinctly different from each other; we found no common ancestors among them, unlike the interspecies phylogenetic analysis where evolutionary positions are already fixed. In this manuscript, the consensus-based method was used (Beura et al. 2023),(Aziz et al. 2022),(Sen et al. 2021), where we derived the consensus sequence by finding the most frequently occurring nucleotide in every position throughout the whole sequence across all strains. This method also proved some fundamental facts of molecular evolution, such as *ti* is greater than *tv*, synonymous SNPs are more than non-synonymous SNPs, and single nucleotide polymorphisms are much more prevalent than double or triple nucleotide polymorphisms. Moreover, this method also proves that when there is a double mutation in a single position of a codon, the combination of mutations is more of a *ti, tv* type rather than a *tv, tv* type.

In a study by Piyali et al.(Sen et al. 2021), the authors reported that the mutation frequency at four-fold degenerate (FFD) sites is higher than in intergenic regions (IRs). This might be because FFD sites are influenced by both replication and transcription, whereas IRs are influenced by replication. Though both IRs and FFD are considered under weak purifying selection, the magnitude of the purifying selection may be different in different regions. In this context, it is interesting that intra-operon intergenic regions, which experience both replication and transcription, exhibit a low frequency of SNPs in comparison to inter-operon intergenic regions that primarily undergo replication alone (Beura et al. 2023). It is argued that stronger selection acting on functionally important sequences within the intra-operon intergenic regions, such as ribosome-binding sites (RBS), might be responsible for the low SNP occurrence at intra-operon intergenic regions. It is pertinent to note that the frequency of SNPs in the intergenic regions is lower than the frequency of SNPs in coding sequences we have studied here. These findings highlight how some SNPs are influenced by transcription but not replication. In conclusion, transcription-induced mutagenesis remains an emerging and important area in molecular evolutionary research, particularly for understanding how DNA repair, oxidative stress, and transcription collectively shape genome evolution.

## Methodology

### Coding sequence information

We analyzed synonymous SNPs in 2091 coding sequences (CDS) by aligning sequences of 157 strains of *E. coli* (Thorpe et al. 2017). We used three criteria to consider a CDS for the SNP study. First, the CDS size remained uniform across the strains; there were no INDELs (insertions and deletions), and there were no ambiguous nucleotides ‘N’,’?’. Further, a CDS with the above three criteria and available from a minimum of 100 out of 157 strains was considered for SNP analysis. Usually, a smaller-sized CDS having a lesser number of SNPs was avoided.

SNPs were analyzed codon-wise as synonymous and non-synonymous. In this study, synonymous SNPs at a single codon position across the strains were considered. It is pertinent to note that polymorphism at two or three positions in a codon was found in less than 5%, which was not of our interest for this study, as our focus was on synonymous SNPs at a single position of a codon.

Our SNP analysis was based on a reference sequence-based approach (Aziz et al. 2022), (Sen et al. 2022), (Beura et al. 2023). In this approach, a reference sequence is generated where the most common nucleotide at each position across the CDS aligned was considered as the reference nucleotide. The method has been demonstrated by taking an example of a hypothetical CDS in Fig. S4. The most frequently occurring nucleotide was assumed to represent the original/ancestral form.

The SNP data were verified using three approaches: strain-wise, phylogeny-wise, and CDS-wise verification. In strain-wise verification, each strain was compared to a reference sequence, and any strain showing significant dissimilarity from the expected threshold was considered an outlier and removed from the dataset. For phylogeny-wise verification, a phylogenetic tree was constructed using all strains, allowing the identification and removal of outliers based on their unusual placement in the tree. In CDS-wise verification, SNP counts were assessed in 100 bp segments (or more) of the CDS sequences; if any segment exhibited an unexpectedly high number of SNPs, the contributing strain(s) were examined and removed from the dataset.

It is pertinent to note that, if a polymorphism arises during an early replication cycle, it will be inherited by a larger proportion of descendant strains, resulting in a higher number of altered nucleotides at that specific genomic position. For example, if we observe a high number of C→T substitutions at the same site across multiple strains, we interpret this as a single mutational event rather than multiple independent occurrences. For instance, in a dataset comprising 157 strains, if 80 strains exhibit a cytosine (C) and 77 exhibit a thymine (T) at a particular genomic position, we consider this to represent a single C→T mutation event. The relatively high number of T bases at this position suggests that the polymorphism occurred early in the replication or evolutionary timeline. Thus, in our analysis, the frequency of T at a given position reflects the timing of the mutational event, not its multiplicity.

### Calculation of synonymous SNPs by considering codon degeneracy

SNPs can result in one nucleotide being substituted by any of the other three, leading to nine possible codon substitutions for each original codon. The synonymous transition (*sti*) and synonymous transversion (*stv*) frequency values were being normalized by considering codon degeneracy (Beura et al.),(Beura 2024).In this study, there are a total of 12 types of SNPs that have been studied, out of which 4 are *ti*s, and 8 are single-nucleotide *tv*s. In every CDS, the SNP is normalized for each type of substitution. Suppose in a CDS we found C→T SNP number as X, but the total possible synonymous C→T number as Y, then the C→T frequency value for that CDS would be / The same process has been followed for quantifying each *ti*s (G→A, A→G, T→C) and each *tvs* (A→C, A→T, C→A, C→G.T→A, T→G, G→C, G→T)

We calculated synonymous mutation frequency for all the selected 2091 coding sequences using a program. The Python script used for SNP analysis is freely available at: https://github.com/CBBILAB/Strand-Asymmetry-SNP-Analysis-Tool

### Distribution of CDS in the LeS and LaS

The 2091 CDS of *E. coli* were assigned into the LeS and the LaS by using the GC skew method. (Lobry 1996). The formula used to calculate the GC skew was 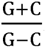. The positive GC skew combined with a “+” strand direction indicates that the CDS is located on the LeS, whereas a positive skew with a “–” strand direction indicates its presence on the LaS. Similarly, a negative GC skew with a “+” strand denotes a lagging-strand CDS, while a negative skew with a “–” strand corresponds to a leading-strand CDS. CDS positions were estimated using the Genome GC Skew diagram (https://atgcgraphview.netlify.app/). The GC skew diagram was shown in Fig. S5. Out of the 2091 selected CDS, 1,166 were located on the LeS, while the remaining 925 were on the LaS. CDS information is provided in Table S4.

### SNP analysis based on CDS size

The filtered 2091 CDS are further classified based on the size of the CDS into the other three categories, CDS size ≥ 500 bp (1775 CDS), ≥ 1000 bp (946 CDS), and ≥ 2000 bp (150 CDS).All CDS mutation frequencies were provided in Table S5. It is thought that CDS of larger size will be more truly representative of the polymorphism spectra, because of the higher abundance of SNPs in larger size CDS. We observed a lesser number of SNPs where CDS below 500 bp (316 CDS) were considered (Table S6)

### Analysis of PR2 in Les and Las CDS

Chargaff’s second parity rule, which is known as PR2, demonstrates that in single-stranded DNA, abundance values of complementary nucleotides are similar (A∼T; G∼T). In this study, we have compared the frequencies of complementary *ti* and complementary *tv* SNPs. If the values are similar, we thought that it maintained the parity rule; if they differ significantly, we thought it violated the PR2. Complementary *ti*, such as C→T and G→A (T→C and A→G), and complementary *tv*, including A→C and T→G, A→T and T→A, C→A and G→T, and C→G and G→C, are quantified in both LeS and LaS CDS to determine if they maintain or violate PR2 It was further confirmed by performing the Mann-Whitney test (https://www.statskingdom.com/170median_mann_whitney.html). To account for minor variations, a threshold was established. Complementary pairs with a difference within 5% of the mean of their average frequencies were classified as equal or borderline. Violation of PR2 often signifies that the asymmetric mutational or selective process is acting on that region related to replication and transcription. The Pearson Chi-Square statistic was also performed to find the strand dependency of complementary SNPs in both LeS and LaS CDS groups (https://www.socscistatistics.com/tests/chisquare/)

### Comparison of SNP frequency between LeS and LaS CDS

The CDS are separated based on their distribution on the LeS or LaS. The frequency of each SNP type was calculated independently for both strands, and the mean frequency values were used for comparative analysis between the two groups. To assess statistical differences between CDS located on the LeS and LaS strands, the Mann–Whitney U test was applied across all size classifications.

### Analysis of the effect of gene expression on strand-specific complementary SNPs using codon adaptation index (CAI) and protein expression data

CDS were ranked based on CAI and protein expression values separately for LeS and LaS. Based on this ranking, two expression groups were defined: CDS with the highest CAI values (highly expressed group) and CDS with the lowest CAI values (lowly expressed group). The analysis was done on CDS with sizes greater than 1000 bp. From each strand, we selected the top 50 CDS with the highest CAI values and the bottom 50 CDS with the lowest CAI values and designated them as highly expressed and lowly expressed genes, respectively. The same experiment was performed using protein expression data, For both LeS and LaS, the disparity in substitution frequencies was compared between the highly expressed genes and the lowly expressed genes groups across all substitution types. The Pearson Chi-Square statistic was also performed to find the expression dependency of complementary SNPs in both LeS and LaS CDS groups (https://www.socscistatistics.com/tests/chisquare/).

## Availability of data and materials

The datasets generated and/or analyzed during the current study are available in the GitHub repository. The source code used for analysis is publicly available at GitHub: https://github.com/CBBILAB/Strand-Asymmetry-SNP-Analysis-Tool.

## Competing Interest Statement

The author(s) declare no conflict of interest

## Author Contributions

ND: conceptualised, methodology, working, analysis, discussion, and writing; PKB: methodology, writing and discussion; PS: methodology, software development, writing, and discussion; R.A: methodology, writing and discussion; AKB: conceptualised, working, and discussion; DK, P.S,MJ, NDN, RCD, EJF.: discussion and writing; SSS, and SKR.: design, critical analysis, writing, discussion, and supervision

## Acknowledgments

ND. is thankful to the DBT, GoI-funded Centre for Bioinformatics and Computational Biology, Tezpur University (BT/PR40253/BTIS/137/52/2022) for the fellowship of Scientific Administrative Assistant. N.D.N, RCD, SSS & SKR thankfully acknowledge the DBT-funded Bioinformatics and Computational Biology Centre at Tezpur University (BT/PR40253/BTIS/137/52/2022). PB, SKR, and SSS fully acknowledge the grant from DBT (NNP Project).

## Notes

### Competing Interest Statement

The authors have declared no competing interest.

### Summary of Updates

Figures updated, Pearson chi-square test included, supplementary files updated, results and discussion section updated

